# A meta-analysis of 3-nitrooxypropanol effects on methane production and yield in beef cattle

**DOI:** 10.1101/2025.08.21.671547

**Authors:** M. H. de Oliveira, R. Zihlmann, A. Bannink, K. A. Beauchemin, J. Dijkstra, E. M. Pressman, S. van Gastelen, E. Kebreab

**Affiliations:** Department of Animal Science, University of California, Davis, CA 95616, USA; DSM Nutritional Products, Animal Nutrition & Health, PO Box 2676, 4002 Basel, Switzerland; Wageningen Livestock Research, Wageningen University & Research, PO Box 338, 6700 AH, Wageningen, the Netherlands; Lethbridge Research and Development Centre, Agriculture and Agri-Food Canada, 5403 1st Avenue South, Lethbridge, Alberta T1J 4B1, Canada; Animal Nutrition Group, Wageningen University & Research, PO Box 338, 6700 AH, Wageningen, the Netherlands

**Keywords:** Feed additive, greenhouse gas, livestock, mathematical modelling, mitigation

## Abstract

Beef cattle are a major source of enteric methane (CH_4_) emissions, a potent greenhouse gas (GHG). The feed additive 3-nitrooxypropanol (3-NOP) has been shown to reduce CH_4_ emissions by inhibiting methyl-coenzyme M reductase, an enzyme critical to methanogenesis in archaea. This study aimed to quantify the effects of 3-NOP on CH_4_ production (g/d) and yield (g/kg DM intake; DMI) in beef cattle and to evaluate how diet composition influences the mitigation response. A systematic literature review identified 17 peer-reviewed in vivo studies, yielding 45 treatment means. Treatment effects were expressed as absolute and relative mean differences versus control groups. Predictor variables included 3-NOP dose, 3-NOP dose^2^, DMI, dietary concentration of NDF, CP, starch, fat, and organic matter (OM), roughage proportion, BW, and dietary inclusion of monensin (yes/no). Four types of models were developed, all including the intercept and 3-NOP dose as fixed predictors, differing as follows: (model 1) optional inclusion of 3-NOP dose^2^ when *P* < 0.10; (model 2) model 1 plus pre-inclusion of NDF concentration; (model 3) pre-inclusion of NDF concentration plus additional predictors (pairwise r ≤ 0.5) that significantly improved model accuracy (*P* < 0.10); and (model 4) additional predictors selected under the same criteria as model 3, without pre-inclusion of NDF concentration. For models 3 and 4, a maximum of 5 predictors were considered and evaluated using leave-one-out cross-validation. Across studies, 3-NOP doses ranged from 32 to 338 mg/kg of DM. On average, 3-NOP reduced CH_4_ production by 49.9 ± 28.61 g/d (36.2 ± 24.42%) and CH_4_ yield by 5.3 ± 3.61 g/kg DMI (33.2 ± 25.54%). The best models were selected based on biological interpretability, statistical significance, and predictive accuracy (as measured by RMSE) and included 3-NOP dose, dietary NDF concentration, DMI, and BW as significant predictors (the latter two only for absolute CH_4_ production). Mitigation efficacy increased with higher DMI and declined with increasing NDF concentration and BW. Absolute reductions of 53.1 g/d and 5.88 g/kg of DMI, and relative reductions of 37.6% in CH_4_ production and 35.0% in CH_4_ yield were predicted when moderators were at their mean value (3-NOP dose = 134.4 mg/kg of DM; NDF concentration = 32.8% of DM; DMI of 8.6 kg/d). These results support the effectiveness of 3-NOP in mitigating enteric CH_4_ emission in beef cattle and provide quantitative models to be used in assessment tools and GHG inventory methodology.

**Implications:** The feed additive 3-nitrooxypropanol effectively reduces enteric methane emissions in beef cattle. This meta-analysis found average reductions of 36.2% in methane production and 33.2% in methane yield. Efficacy depended on diet composition; declining with increasing NDF concentration for both methane production (g/d) and yield (g/kg of DM intake; DMI). Greater DMI increased absolute methane production reduction but did not influence absolute methane yield reduction or relative reduction of both methane production and yield. These results support the targeted use of 3-nitrooxypropanol as a mitigation strategy and provide empirical models to inform greenhouse gas inventories and carbon accounting.

## Introduction

Ruminant livestock systems are a major contributor to global anthropogenic methane (CH_4_) production, accounting for 12% of total global greenhouse gas (GHG) emissions (FAO, 2022; 2015 estimates). Within the livestock sector, enteric CH_4_ contributes approximately 78% of total CH_4_ emissions (FAO, 2022), making it a critical target for mitigation strategies. In beef cattle systems, average enteric CH_4_ emissions have been estimated at 161 g/animal per day (van Lingen et al., 2019), contributing to the environmental footprint of beef production.

Addressing enteric CH_4_ emissions has become a priority for sustainable livestock production, spurring research into nutritional interventions that can reduce methanogenesis without impairing animal performance and health. Among these interventions, 3-nitrooxypropanol (3-NOP) has emerged as one of the most effective and well-characterized additives (Belanche et al., 2025). The 3-NOP acts as a structural analog of methyl-coenzyme M, selectively inhibiting the enzyme methyl-coenzyme M reductase (MCR) in methanogenic archaea (Duin et al., 2016). This targeted mechanism is selective for methanogens only and minimizes disruptions to rumen function while achieving significant CH_4_ reduction. Beyond its anti-methanogenic properties, 3-NOP supplementation has been associated with improvements in feed efficiency and ruminal fermentation patterns in beef cattle (Orzuna-Orzuna et al., 2024), with no adverse effects on animal performance (Kim et al., 2019) or indications of mutagenic or genotoxic risk (Thiel et al., 2019).

The efficacy of 3-NOP has been demonstrated across multiple studies, with reported mitigation rates varying widely depending on dose, animal category, and dietary context. Previous meta-analysis by Dijkstra et al. (2018) found that CH_4_ production was reduced by 22.2% in beef cattle and 39.0% in dairy cattle at a mean 3-NOP dose of 123 mg/kg of DM and a mean NDF concentration of 33.1% DM. That analysis, however, was limited to only six beef and five dairy studies, reducing its ability to capture broader patterns in mitigation outcomes. Since then, the number of in vivo trials evaluating 3-NOP supplementation has increased, enabling further meta-analytical assessments. Five recent meta-analyses have examined the CH_4_ mitigating effects of 3-NOP in ruminants. These studies, however, either focused exclusively on dairy cattle (Kebreab et al., 2023; Martins et al., 2024), did not investigate the causes of heterogeneity in the response to 3-NOP supplementation in beef cattle using multivariate approaches (Almeida et al., 2021; Orzuna-Orzuna et al., 2024), or combined beef and dairy data (Macdonald et al., 2024), reducing predictive accuracy for specific production system. Besides that, the latter study presented equations with contradictory NDF effects that vary between positive and negative associations along with extreme predictions even within normal ranges of predictor values. Therefore, a robust quantitative synthesis exclusively from beef cattle systems remains necessary to develop predictive models and to explain how diet composition and 3-NOP dose level affects CH_4_ mitigation. These predictions may be applied to optimize the use of 3-NOP across diverse production systems and to support policy and management decisions aimed at improving sustainability in the livestock sector (Dijkstra et al., 2025).

The objective of this study was to quantitatively assess the sources of variability in CH_4_ mitigation response in beef cattle using a meta-analytical approach. A database was synthesized to 1. quantify the efficacy of 3-NOP in reducing CH_4_ production and yield; 2. evaluate how dietary factors modulate the effect of 3-NOP; and 3. develop predictive models that inform about the efficacy of 3-NOP supplementation. We hypothesized that 3-NOP supplementation reduces enteric CH_4_production (g/d) and yield (g/kg DM intake; DMI) in beef cattle, and that the magnitude of this reduction increases nonlinearly, approaching a plateau at elevated 3-NOP doses. Additionally, we hypothesized that higher dietary NDF concentration diminishes the 3-NOP mitigation response due to altered rumen fermentation dynamics.

## Materials and Methods

### Data collection

A comprehensive systematic search was performed using Web of Science (Thomson Reuters Science, https://www.webofscience.com/), Scopus (Elsevier, https://www.scopus.com/), and Google Scholar (https://scholar.google.com/) based on the keywords “3-NOP” (including all variants, such as “nitrooxypropanol” and the commercial name “Bovaer”) + “beef cattle”. To be included, studies had to meet the following criteria: (1) report in vivo trials with beef cattle, (2) include a control group without 3-NOP supplementation, (3) present CH_4_ emission data as g/d or g/kg of DMI, and (4) deliver 3-NOP through the diet. A full description of the search process is provided in Figure 1. Among the databases, 256 studies were obtained. After removing 117 duplicates, 139 unique studies were screened. Of these, 121 were excluded for not meeting the inclusion criteria as following: occasional wrong population (e.g., dairy, lambs; n = 48), reviews (n = 41), rumen microbiome or in vitro trials (n = 10), meta-analyses (n = 8), whole-farm, life cycle assessments or modeling approaches (n = 6), manure assessments (n = 4), toxicity studies (n = 2), behavior (n = 1), and mechanisms (n = 1). Eighteen studies were assessed for eligibility, and one was rejected (McGinn et al., 2019) because it did not measure absolute CH_4_ emissions.

**Fig. 1.**
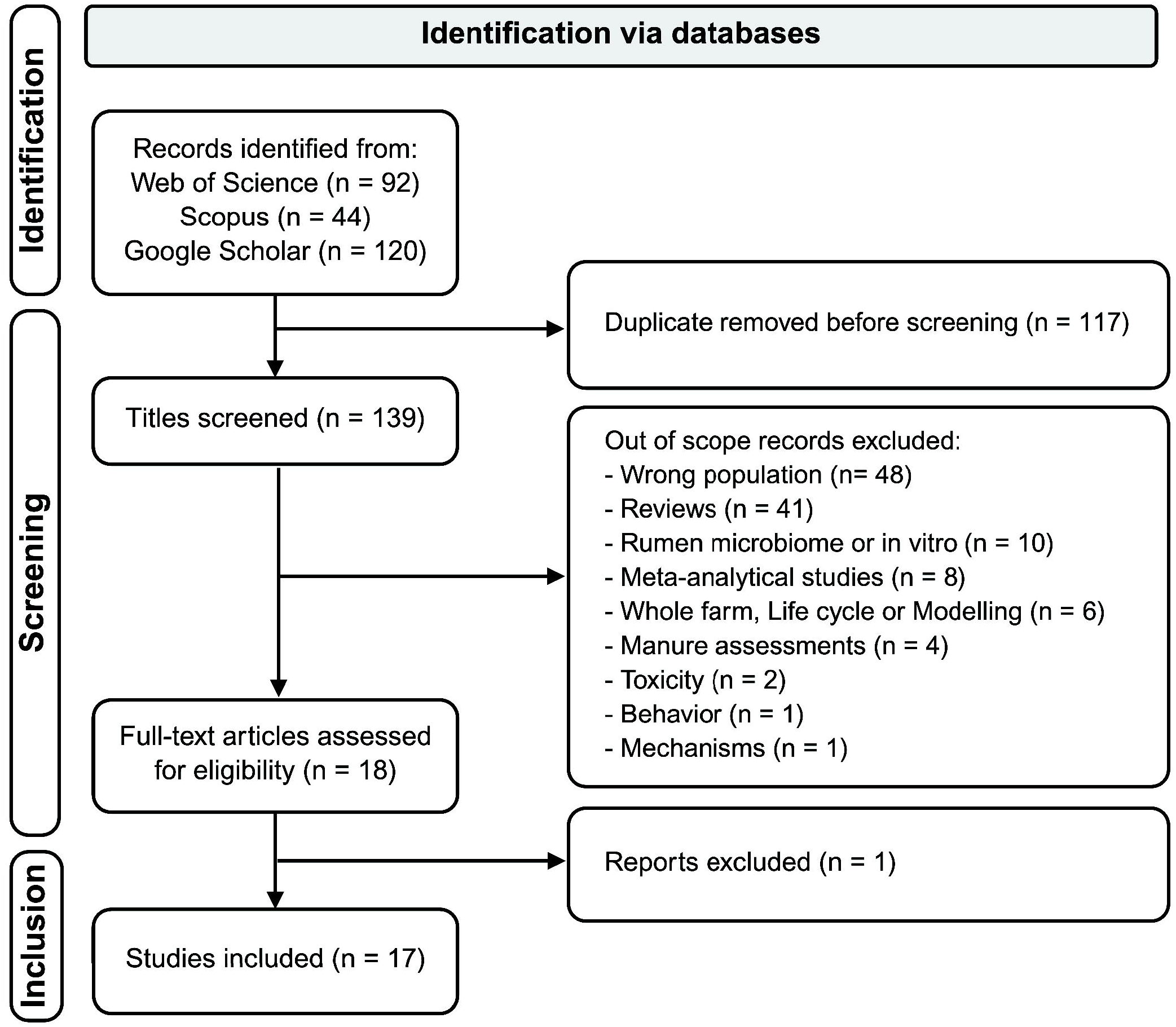
Flowchart showing the search strategy and eligible studies for the meta-analysis on the methane mitigating effects of 3-NOP (3-nitrooxypropanol) in beef cattle diets. 850

Seventeen studies met the inclusion criteria and were included in the meta-analysis, providing a total of 45 treatment means (Table 1; Romero-Perez et al., 2014; Romero-Perez et al., 2015; Vyas et al., 2016; Vyas et al., 2018a; Vyas et al., 2018b; Martinez-Fernandez et al., 2018; Kim et al., 2019; Alemu et al., 2021a; Alemu et al., 2021b; Zhang et al., 2021; Alemu et al., 2023; Almeida et al., 2023; Araújo et al., 2023; Kirwan et al., 2024; Pedrini et al., 2024; Souza et al., 2024; Burgers et al., 2025). Fourteen studies reported the specific production phase of beef cattle system (Backgrounding/Growing or Finishing). For the three studies lacking this information (Martinez-Fernandez et al., 2018; Zhang et al., 2021; Alemu et al., 2023), the production phase was inferred based on dietary composition, classifying them as backgrounding trials due to the high-forage diets fed (≥ 90% forage on DM basis). Methane emissions were quantified using the GreenFeed system (C-Lock Inc.; 7 experiments), respiration chambers (9 experiments), or the sulfur hexafluoride (SF_6_) tracer gas technique (1 experiment), which are accepted methods for measuring enteric CH_4_ production in ruminants (Hristov et al., 2013; Hammond et al., 2016; Hristov et al., 2025).

**Table 1.**
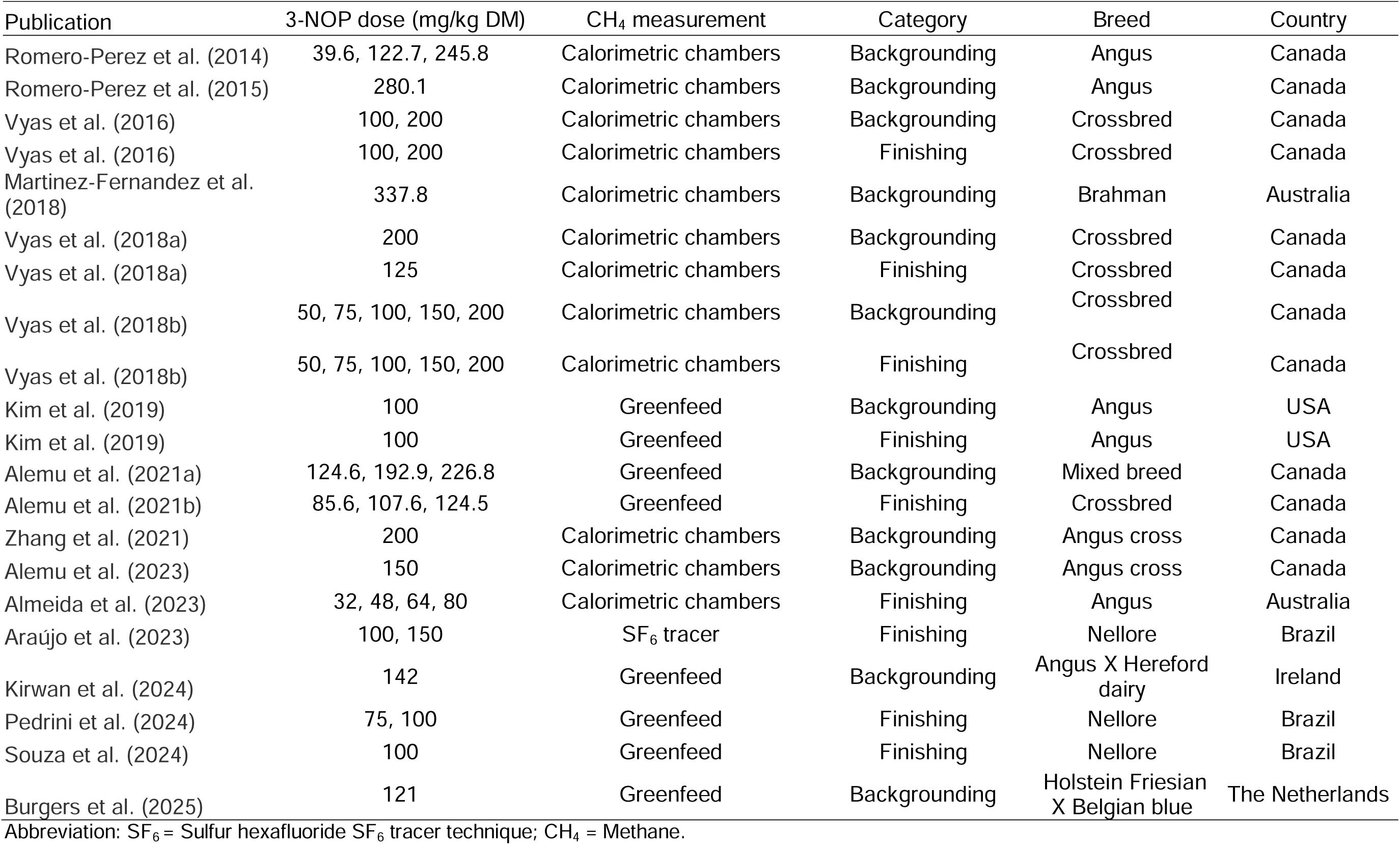
Format tables. Description of studies included in the meta-analysis database

### Response Variables

The key response variables examined were CH_4_ production (g/d) and CH_4_ yield (g/kg of DMI); CH_4_ intensity (g/kg of BW; g/kg of average daily gain; ADG; or g/kg of carcass ADG) was not included as it was not reported in most of the selected studies. The selected studies presented both response variables separated by treatment group as least-squares means with a common pooled standard error. Treatment contrasts were calculated as: 1. mean difference (MD), calculated as:

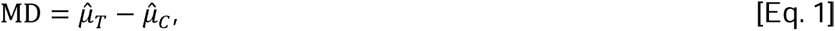

Where 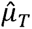 and 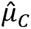 represent the reported least-squares means for the treatment and control groups, respectively; 2. relative mean difference (RMD), expressed as a percentage and calculated as:

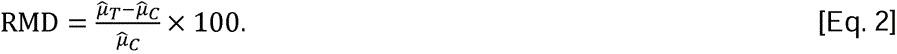

If variances of contrasts were not explicitly reported, the variance of MD was calculated assuming independence between the least-squares means:

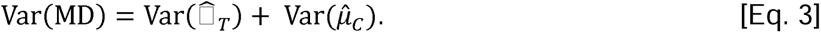

The variance of RMD was approximated using the delta method (Franz, 2007) with the following formula:

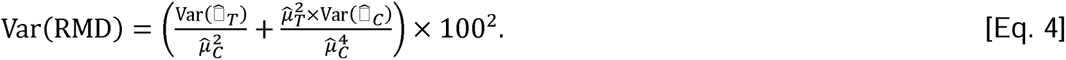

These variances were then used to compute inverse-variance weights, assigning greater influence on comparisons with higher precision (i.e., lower variance).

### Explanatory Variables

Potential explanatory variables were extracted from the studies. These included 3-NOP dose (mg/kg of DM), DMI (kg/d), dietary concentrations of NDF (% of DM), crude fat (% of DM), CP (% of DM), starch (% of DM), and organic matter (OM; % of DM), roughage proportion (% of diet DM), BW (kg), and inclusion of monensin (yes/no).

Several studies did not report complete dietary composition data, particularly for starch, fat, or OM. Missing values were estimated using the approach described by Kebreab et al. (2023). Individual feed ingredient compositions were obtained from NASEM (2021) or Feedipedia (https://www.feedipedia.org) when not available in NASEM (2021). These values were weighted by their inclusion rate in the diet to estimate overall dietary composition. Calculated values showed close agreement with reported values, with average relative deviations of 10.5% for starch, -3% for fat, and 0.5% for OM. A summary of the database is provided in Table 2.

**Table 2.**
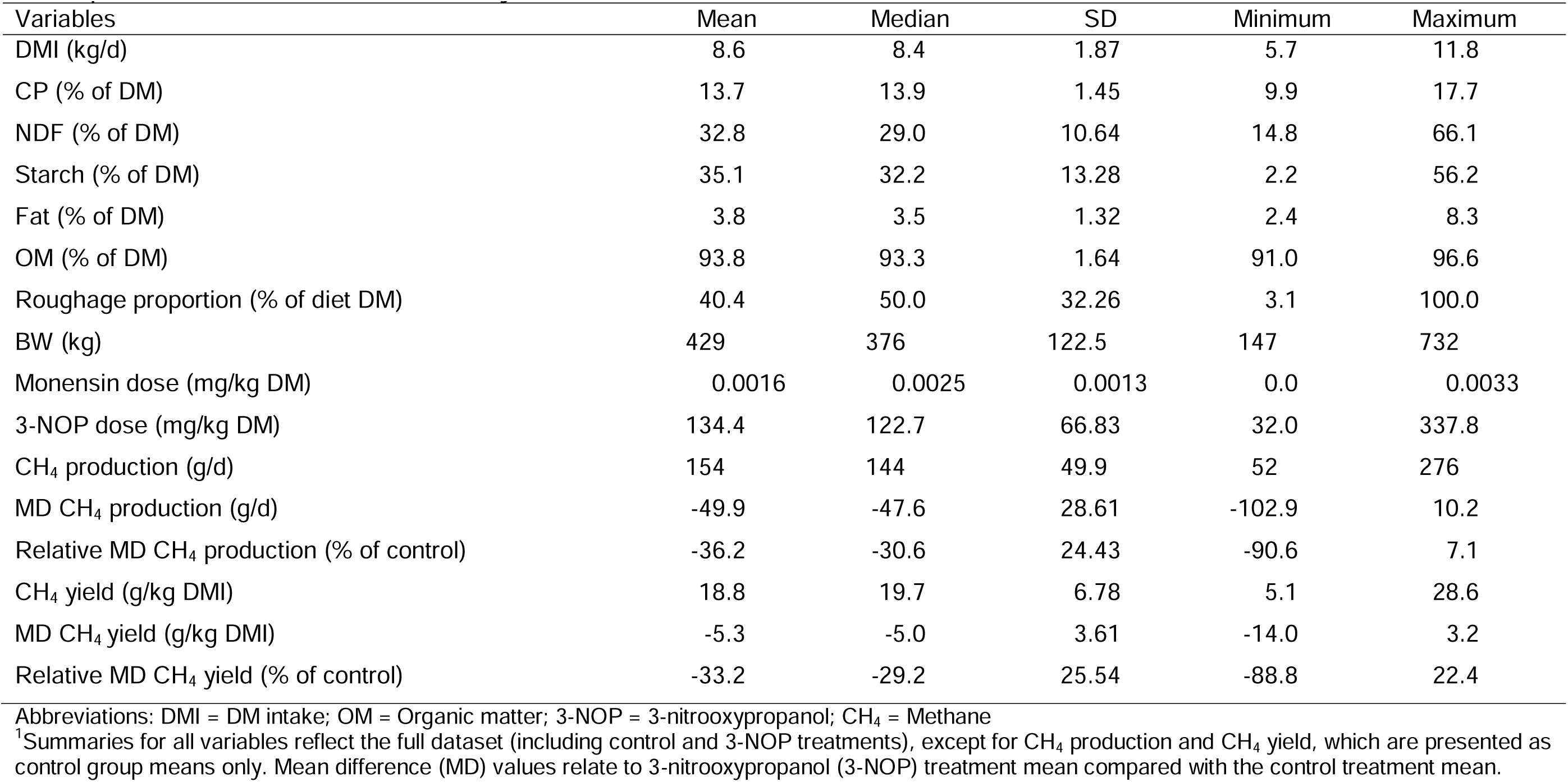
Descriptive statistics of feed intake, dietary characteristics, and methane emission^1^.

| Variables | Mean | Median | SD | Minimum | Maximum |
| --- | --- | --- | --- | --- | --- |
| DMI (kg/d) | 8.6 | 8.4 | 1.87 | 5.7 | 11.8 |
| CP (% of DM) | 13.7 | 13.9 | 1.45 | 9.9 | 17.7 |
| NDF (% of DM) | 32.8 | 29.0 | 10.64 | 14.8 | 66.1 |
| Starch (% of DM) | 35.1 | 32.2 | 13.28 | 2.2 | 56.2 |
| Fat (% of DM) | 3.8 | 3.5 | 1.32 | 2.4 | 8.3 |
| OM (% of DM) | 93.8 | 93.3 | 1.64 | 91.0 | 96.6 |
| Roughage proportion (% of diet DM) | 40.4 | 50.0 | 32.26 | 3.1 | 100.0 |
| BW (kg) | 429 | 376 | 122.5 | 147 | 732 |
| Monensin dose (mg/kg DM) | 0.0016 | 0.0025 | 0.0013 | 0.0 | 0.0033 |
| 3-NOP dose (mg/kg DM) | 134.4 | 122.7 | 66.83 | 32.0 | 337.8 |
| CH<sub>4</sub> production (g/d) | 154 | 144 | 49.9 | 52 | 276 |
| MD CH<sub>4</sub> production (g/d) | -49.9 | -47.6 | 28.61 | -102.9 | 10.2 |
| Relative MD CH<sub>4</sub> production (% of control) | -36.2 | -30.6 | 24.43 | -90.6 | 7.1 |
| CH<sub>4</sub> yield (g/kg DMI) | 18.8 | 19.7 | 6.78 | 5.1 | 28.6 |
| MD CH<sub>4</sub> yield (g/kg DMI) | -5.3 | -5.0 | 3.61 | -14.0 | 3.2 |
| Relative MD CH<sub>4</sub> yield (% of control) | -33.2 | -29.2 | 25.54 | -88.8 | 22.4 |

Abbreviations: DMI = DM intake; OM = Organic matter; 3-NOP = 3-nitrooxypropanol; CH<sub>4</sub> = Methane
<sup>1</sup>Summaries for all variables reflect the full dataset (including control and 3-NOP treatments), except for CH<sub>4</sub> production and CH<sub>4</sub> yield, which are presented as control group means only. Mean difference (MD) values relate to 3-nitrooxypropanol (3-NOP) treatment mean compared with the control treatment mean.

### Statistical Analyses

#### Model development, inspection and selection

All statistical analyses were performed using R version 4.4.3 (R Core Team, 2024) with the “metafor” package (version 4.6.0; Viechtbauer, 2010). Mixed-effect meta-regression models (Supplementary Material S1) were constructed by incorporating different combinations of fixed predictors. Because multiple treatment groups within individual studies often shared a common control, we used robust variance estimation (Hedges et al., 1999) to account for the non-independence of treatment comparisons within studies.

The pool of potential fixed effects included the 10 explanatory variables previously described (3-NOP dose, DMI, dietary contents of NDF, crude fat, CP, starch, and OM, roughage proportion, BW, and inclusion of monensin), plus an additional quadratic term for 3-NOP dose to evaluate potential nonlinear responses, bringing the total to 11 candidate predictor variables, all of which (except monensin inclusion being a class variable) were centered on their means before analysis. Model selection was conducted across four procedures. Based on previous evidence as a key dietary factor influencing 3-NOP efficacy (Dijkstra et al., 2018; Kebreab et al., 2023), NDF concentration was pre-included in two of the procedures. Briefly, modeling consisted of the following approaches, all including the overall mean (intercept) and 3-NOP dose as fixed predictors, plus:

Model 1) a quadratic term for 3-NOP dose (3-NOP dose^2^) if statistically significant (*P* < 0.10).

Model 2) NDF concentration and 3-NOP dose^2^, with the latter considered only if statistically significant (*P* < 0.10).

Model 3) NDF concentration, allowing additional predictors with pairwise correlation ≤ 0.5 and that significantly (*P* < 0.10) improved prediction accuracy.

Model 4) Any additional predictor with pairwise correlation ≤ 0.5 that significantly (*P* < 0.10) improved model accuracy.

For models 3 and 4, up to five predictors (excluding the intercept) were considered. Inclusion of production phase (backgrounding vs. finishing) as a categorical moderator was also evaluated, but because this yielded similar prediction results, the data were pooled in the final analysis to increase model robustness and statistical power.

Model performance was assessed based on RMSE derived from leave-one-out cross-validation (LOOCV). The LOOCV was implemented at the study level, meaning that all treatment comparisons from a given publication were simultaneously excluded from the training data. This approach provides a more realistic estimate of prediction error for future studies compared to validation at the individual treatment comparison level. Pseudo R^2^ values were calculated to estimate the proportion of between-study heterogeneity accounted for by the included moderators (Raudenbush, 2009). Total heterogeneity was quantified using the *Ι* ^2^ statistic, representing the proportion of total variability attributable to true heterogeneity rather than sampling error relative to the total amount of variance. Then, it was decomposed into between-trial heterogeneity (*Ι* ^2^_1_) representing variability across different publications, and between-comparison heterogeneity (*Ι* ^2^_2_) representing variability among treatment contrasts within trials, both relative to the total amount of variance. The degree of heterogeneity based on *Ι* ^2^ statistic was classified as follows: low (0– 25%), moderate (25–50%), substantial (50–75%), and high (>75%) (Higgins et al., 2003). The Q-test was used to assess the statistical significance of heterogeneity.

Following model selection, diagnostic checks were conducted to verify model assumptions. Residual plots were examined for patterns suggesting heteroscedasticity or nonlinearity. Cook’s distance values were calculated to identify potentially influential observations (Cook, 1977). Potential publication bias was assessed through visual inspection of funnel plots and complementary statistical tests, including Egger’s test (Egger et al., 1997).

## Results

### Study characteristics

Seventeen studies published between 2014 and 2025 were included in the meta-analysis (Table 1). These studies covered backgrounding and finishing phases of beef production and involved *Bos taurus* and *Bos indicus* cattle across six countries, including Canada (n = 9), Brazil (n = 3), Australia (n = 2), Ireland (n = 1), the Netherlands (n = 1), and USA (n = 1). Descriptive statistics for dietary and animal variables are summarized in Table 2. Average BW was 429 ± 122.5 kg. The mean DMI was 8.6 kg/d, ranging from 5.7 to 11.8 kg/d. The dietary NDF concentration averaged 32.8%, and average 3-NOP dose was 134.4 mg/kg DM. Mean CH_4_ production and yield in control groups were 154 g/d and 18.8 g/kg DM, respectively.

Study-level effects of 3-NOP supplementation on CH_4_ production and yield, stratified by production phase are presented in Figure 2. Relative MD to control was mostly negative across studies included in the analysis. The magnitude of the reduction was generally larger in finishing cattle than in backgrounding cattle. Although a few studies reported minimal or slightly positive changes, the majority showed reductions in both CH_4_ production and yield regardless of production phase.

**Fig. 2.**
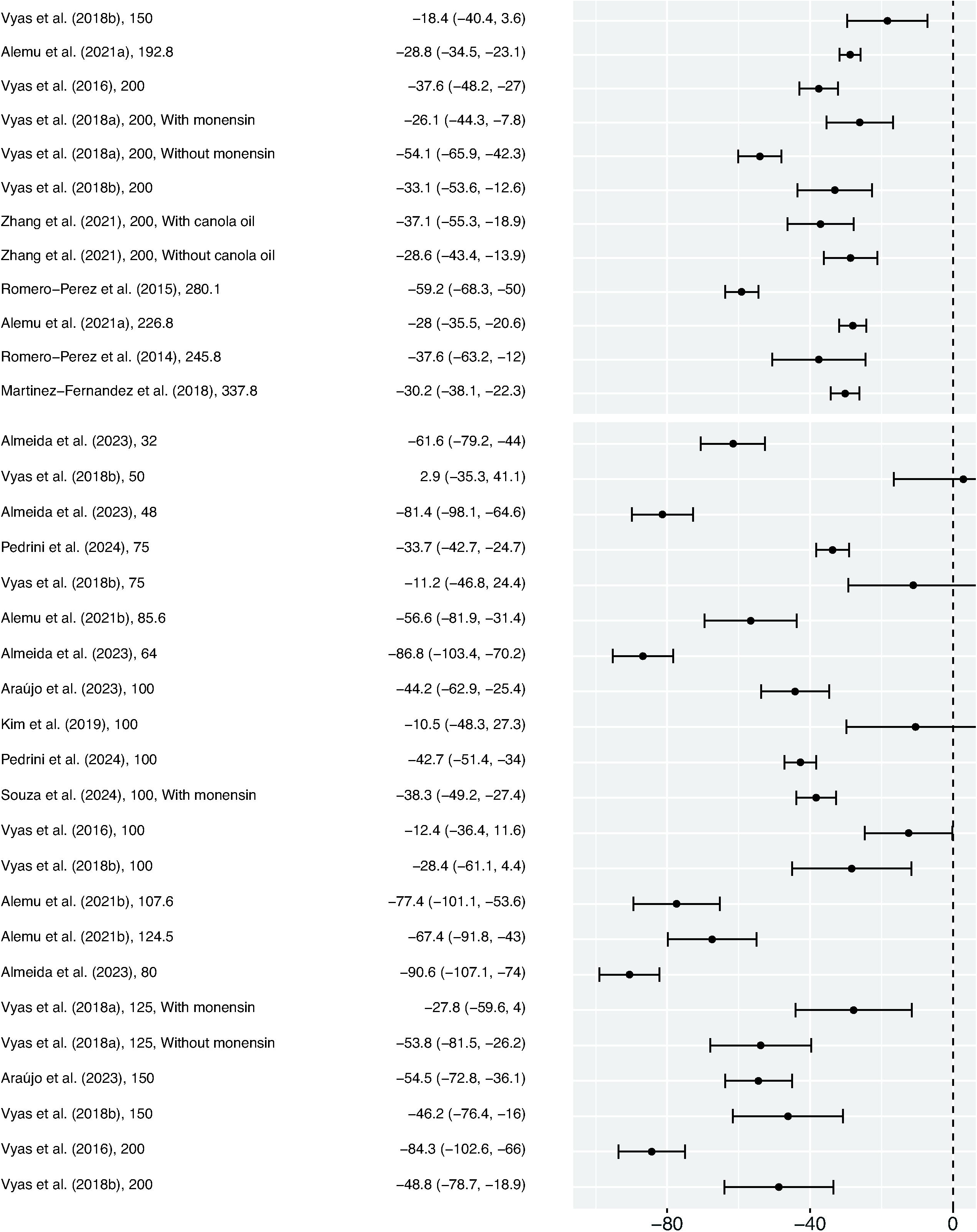
Forest plots of relative mean difference to control in CH_4_ production (g/d) and CH_4_ yield (g/kg DM). Abbreviations: 3-NOP = 3-nitrooxypropanol; 95% CI: 95% confidence interval. Abbreviation: CH_4_ = Methane; CH_4_ = Methane; 3-NOP = 3-nitrooxypropanol.

### Mean difference models

The effects of explanatory variables of the four candidate models on MD of CH_4_ production are presented in Table 3. The baseline model 1, which included only the linear and quadratic 3-NOP dose terms, showed a reduction in CH_4_ production with increasing dose (*P* < 0.001), along with a positive quadratic effect (*P* = 0.035). At the mean 3-NOP dose (134.4 mg/kg of DM (Table 2); linear and quadratic term), CH_4_ production was reduced by 57.9 g/d (intercept; *P* < 0.001). This model presented an RMSE of 27.6 g/d and a pseudo R^2^ of 24.4%, with 75% of total *Ι* ^2^, entirely attributed to between-trial heterogeneity (*Ι* ^2^_1_ = 75%). The predicted MD in CH_4_ production (based on Model 1; Table 3) across different 3-NOP doses is illustrated in Figure 3. As 3-NOP dose increases CH_4_ production decreases (0.309 g/d for each 1 mg/kg of DM increase in 3-NOP dose) with smaller reductions at higher doses (>200 mg/kg of DM), where the saturable dose-response relationship becomes apparent. In model 2, the addition of NDF concentration improved model performance (RMSE = 23.5 g/d; pseudo R^2^ = 60.7%), and reduced heterogeneity (*Ι* ^2^ = 63% attributed entirely to *Ι* ^2^_1_) compared to model 1. This model showed a positive association between NDF concentration (*P* = 0.004) and change in CH_4_ production, with an estimated increase of 1.52 g/d of the (negative) value of the MD in CH_4_ production per 1 percentage point increase in NDF concentration. The predicted reduction in CH_4_ production when all predictors are at their mean values was 54.9 g/d (intercept; *P* < 0.001). Models 3 and 4 yielded identical results and identified DMI (*P* = 0.007) and BW (*P* = 0.015) as additional significant predictors, with change in CH_4_ production decreasing by -4.66 g/d per kg increase in DMI. This model also included NDF concentration (*P* = 0.002), 3-NOP dose (*P* < 0.001) as predictors, with lower RMSE (20.0 g/d), higher pseudo R^2^ (71.5%) and lower *Ι* ^2^ (52%) (*Ι* ^2^_1_= 52%) compared to models 1 and 2. When all three predictors are at their mean values, the estimated CH_4_ production reduction was 53.1 g/d (intercept; *P* < 0.001).

**Fig. 3.**
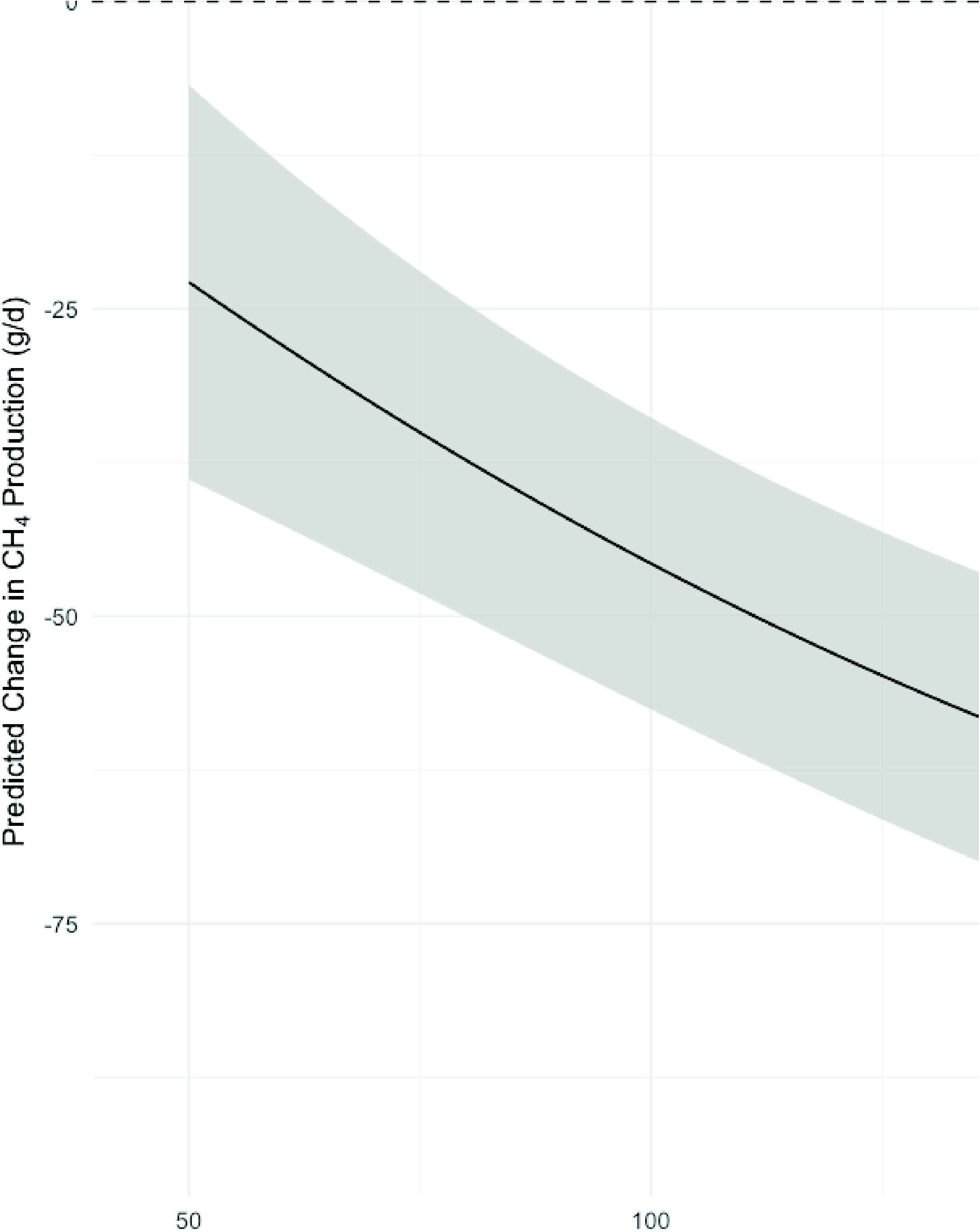
Linear and quadratic effect of 3-NOP dose (mg/kg DM) on CH_4_ production (g/d) reduction in beef cattle. The gray shade area represents the pointwise 95% confidence band. Abbreviation: CH_4_ = Methane; CH_4_ = Methane; 3-NOP = 3-nitrooxypropanol.

**Table 3.** Estimates of overall 3-nitrooxypropanol (3-NOP) effect size and of explanatory variables from models for mean difference in methane production (g/d)

| Variable<sup>1</sup> | Candidate Models<sup>2</sup> |  |  |  |  |  |  |  |  |  |  |  |
| --- | --- | --- | --- | --- | --- | --- | --- | --- | --- | --- | --- | --- |
|  | Model 1 |  |  | Model 2 |  |  | Model 3 |  |  | Model 4 |  |  |
|  | Estimate | SE | P-value | Estimate | SE | P-value | Estimate | SE | P-value | Estimate | SE | P-value |
| DMI (kg/d) |  |  |  |  |  |  | -4.66 | 1.547 | 0.007 |  |  |  |
| NDF (% of DM) |  |  |  | 1.52 | 0.478 | 0.004 | 1.36 | 0.385 | 0.002 |  |  |  |
| BW (kg) |  |  |  |  |  |  | 0.055 | 0.0207 | 0.015 |  |  |  |
| 3-NOP dose (mg/kg DM) | -0.309 | 0.0654 | <0.001 | -0.314 | 0.0403 | <0.001 | -0.352 | 0.0455 | <0.001 |  |  |  |
| 3-NOP dose<sup>2</sup> (mg/kg DM) | 0.0012 | 0.0006 | 0.035 |  |  |  |  |  |  |  |  |  |
| Intercept | -57.9 | 5.04 | <0.001 | -54.9 | 3.79 | <0.001 | -53.1 | 3.22 | <0.001 |  |  |  |
| RMSE (g/d) | 27.6 |  |  | 23.5 |  |  | 20.0 |  |  |  |  |  |
| Pseudo R<sup>2</sup> | 24.4 |  |  | 60.7 |  |  | 71.5 |  |  |  |  |  |
| $I^2$ | 75 | | | 63 | | | 52 | | | | | |
| $I^2_1$ | 75 | | | 63 | | | 52 | | | | | |
| $I^2_2$ | 0 | | | 0 | | | 0 | | | | | |

Abbreviations: DMI = DM intake; 3-NOP = 3-nitrooxypropanol; $I^2$ = percentage of total variability due to heterogeneity; $I^2_1$ = relative between-trial heterogeneity; $I^2_2$ = relative between-comparison heterogeneity.
<sup>1</sup>All variables were centered to their mean value presented in Table 2.
<sup>2</sup>Model 1 = Model with 3-NOP dose as fixed predictor; 3-NOP dose<sup>2</sup> was tested and included only if statistically significant ( $P < 0.10$ ). Model 2 = Model with 3-NOP dose and NDF as fixed predictors; 3-NOP dose<sup>2</sup> was tested and included only if statistically significant ( $P < 0.10$ ). Model 3 = same as model 2, allowing additional predictors with pairwise correlation $\leq 0.5$ that significantly ( $P < 0.10$ ) improved model fit. Model 4 = same as model 1, allowing additional predictors with pairwise correlation $\leq 0.5$ that significantly ( $P < 0.10$ ) improved model fit.
<sup>3</sup>Model 4 is identical to model 3.

For MD of CH_4_ yield (Table 4), models 1 and 4 were identical, including only 3-NOP dose as moderator. The estimated reduction at the mean dose was 5.74 g CH_4_/kg DMI (intercept; *P* < 0.001), with a linear reduction of 0.031 g CH_4_/kg DMI per mg/kg of 3-NOP (*P* < 0.001). This baseline model presented RMSE of 2.9 g/kg DMI and pseudo R^2^ of 45.8%, with 68% of heterogeneity entirely attributed to between-trial heterogeneity (*Ι* ^2^_1_ = 68%). Models 2 and 3, which included NDF concentration (*P* = 0.146) and 3-NOP dose (*P* < 0.001), were identical and estimated a reduction of 5.88 g CH_4_/kg DMI (intercept; *P* < 0.001), when both predictors were at their means.

**Table 4.** Estimates of overall 3-nitrooxypropanol (3-NOP) effect size and of explanatory variables from models for mean difference in methane yield (g/kg DMI)

| Variable<sup>1</sup> | Candidate Models<sup>2</sup> |  |  |  |  |  |  |  |  |  |  |  |
| --- | --- | --- | --- | --- | --- | --- | --- | --- | --- | --- | --- | --- |
|  | Model 1 |  |  | Model 2 |  |  | Model 3 |  |  | Model 4 |  |  |
|  | Estimate | SE | P-value | Estimate | SE | P-value | Estimate | SE | P-value | Estimate | SE | P-value |
| NDF (% of DM) |  |  |  | 0.07 | 0.049 | 0.146 |  |  |  |  |  |  |
| 3-NOP dose (mg/kg DM) | -0.031 | 0.0076 | <0.001 | -0.036 | 0.0076 | <0.001 |  |  |  |  |  |  |
| Intercept | -5.74 | 0.421 | <0.001 | -5.88 | 0.389 | <0.001 |  |  |  |  |  |  |
| RMSE (g/d) | 2.9 |  |  | 2.8 |  |  |  |  |  |  |  |  |
| Pseudo R<sup>2</sup> | 45.8 |  |  | 60.0 |  |  |  |  |  |  |  |  |
| $I^2$ | 68 | | | 61 | | | | | | | | |
| $I^2_1$ | 68 | | | 61 | | | | | | | | |
| $I^2_2$ | 0 | | | 0 | | | | | | | | |

Abbreviations: 3-NOP = 3-nitrooxypropanol; $I^2$ = percentage of total variability due to heterogeneity; $I^2_1$ = relative between-trial heterogeneity; $I^2_2$ = relative between-comparison heterogeneity.
<sup>1</sup>All variables were centered to their mean value presented in Table 2.
<sup>2</sup>Model 1 = Model with 3-NOP dose as fixed predictor; 3-NOP dose<sup>2</sup> was tested and included only if statistically significant ( $P < 0.10$ ). Model 2 = Model with 3-NOP dose and NDF concentration as fixed predictors; 3-NOP dose<sup>2</sup> was tested and included only if statistically significant ( $P < 0.10$ ). Model 3 = same as model 2, allowing additional predictors with pairwise correlation $\leq 0.5$ that significantly ( $P < 0.10$ ) improved model fit. Model 4 = same as model 1, allowing additional predictors with pairwise correlation $\leq 0.5$ that significantly ( $P < 0.10$ ) improved model fit.
<sup>3</sup>Model 3 is identical to model 2.
<sup>4</sup>Model 4 is identical to model 1.

Although NDF concentration was not significantly associated with CH_4_ yield, its inclusion resulted in a modest increase in pseudo R^2^ to 60.0%, with similar RMSE (2.8 g/kg DMI) compared to models 1 and 4. Additionally, residual heterogeneity reduced to 61% and was entirely attributable to between-trial variation (*Ι* ^2^_1_).

### Relative mean difference models

Candidate models of RMD in CH_4_ production are presented in Table 5. Model 1 included only linear 3-NOP dose term (*P* = 0.008), predicting a 34.6% (intercept; *P* < 0.001) reduction at the mean dose, but explained little of the variance (pseudo R^2^ = 0%; RMSE = 27.5%). Model 2 and 3 (identical) improved performance with the inclusion of NDF concentration (*P* < 0.001) compared to model 1, predicting 1.32 percentage point increase in RMD of CH_4_ production for every 1 percentage point increase in NDF concentration. This model also included 3-NOP dose (*P* < 0.001) and predicted a 37.6% reduction (intercept; *P* < 0.001) at mean values, explaining more of the variation (pseudo R^2^ = 34.2%; RMSE = 23.3%). Model 4, which allowed additional predictors beyond 3-NOP dose (*P* = 0.003), selected starch and fat concentration and BW. Starch and fat concentration reduced RMD in CH_4_ production (*P* = 0.001 and *P* = 0.017), whereas it increased with BW (*P* = 0.053). This model predicted a 38.5% reduction (intercept; *P* < 0.001) at mean values and presented the best fit (RMSE = 20.4%; pseudo R^2^ = 60.5%) compared to the former models. Residual heterogeneity ranged from 82 to 95% across all models of RMD in CH_4_ production, primarily due to between-trial variation (*Ι* ^2^_1_ = 73 – 84%).

**Table 5.** Estimates of overall 3-nitrooxypropanol (3-NOP) effect size and of explanatory variables from models for relative mean difference in methane production (%)

| Variable<sup>1</sup> | Candidate Models<sup>2</sup> |  |  |  |  |  |  |  |  |  |  |  |
| --- | --- | --- | --- | --- | --- | --- | --- | --- | --- | --- | --- | --- |
|  | Model 1 |  |  | Model 2 |  |  | Model 3 |  |  | Model 4 |  |  |
|  | Estimate | SE | P-value | Estimate | SE | P-value | Estimate | SE | P-value | Estimate | SE | P-value |
| NDF (% of DM) |  |  |  | 1.32 | 0.311 | <0.001 |  |  |  |  |  |  |
| Starch (% of DM) |  |  |  |  |  |  |  |  |  | -1.03 | 0.276 | 0.001 |
| Fat (% of DM) |  |  |  |  |  |  |  |  |  | -6.94 | 2.664 | 0.017 |
| BW (kg) |  |  |  |  |  |  |  |  |  | 0.049 | 0.0238 | 0.053 |
| 3-NOP dose (mg/kg DM) | -0.167 | 0.0578 | 0.008 | -0.210 | 0.0410 | <0.001 |  |  |  | -0.212 | 0.0628 | 0.003 |
| Intercept | -34.6 | 5.67 | <0.001 | -37.6 | 4.52 | <0.001 |  |  |  | -38.5 | 3.50 | <0.001 |
| RMSE (%) | 27.5 |  |  | 23.3 |  |  |  |  |  | 20.4 |  |  |
| Pseudo R<sup>2</sup> | 0.0 |  |  | 34.2 |  |  |  |  |  | 60.5 |  |  |
| $I^2$ | 95 | | | 90 | | | | | | 82 | | |
| $I^2_1$ | 84 | | | 84 | | | | | | 73 | | |
| $I^2_2$ | 11 | | | 6 | | | | | | 9 | | |

Abbreviations: 3-NOP = 3-nitrooxypropanol; $I^2$ = percentage of total variability due to heterogeneity; $I^2_1$ = relative between-trial heterogeneity; $I^2_2$ = relative between-comparison heterogeneity.
<sup>1</sup>All variables were centered to their mean value presented in Table 2.
<sup>2</sup>Model 1 = Model with 3-NOP dose as fixed predictor; 3-NOP dose<sup>2</sup> was tested and included only if statistically significant ( $P < 0.10$ ). Model 2 = Model with 3-NOP dose and NDF concentration as fixed predictors; 3-NOP dose<sup>2</sup> was tested and included only if statistically significant ( $P < 0.10$ ). Model 3 = same as model 2, allowing additional predictors with pairwise correlation $\leq 0.5$ that significantly ( $P < 0.10$ ) improved model fit. Model 4 = same as model 1, allowing additional predictors with pairwise correlation $\leq 0.5$ that significantly ( $P < 0.10$ ) improved model fit.
<sup>3</sup>Model 3 is identical to model 2.

For RMD in CH_4_ yield (Table 6), model 1 included only a linear 3-NOP dose term (*P* = 0.008). At the mean 3-NOP dose (134.4 mg/kg DM; Table 2), the predicted reduction was 32.2% (intercept; *P* < 0.001). The model explained little of the variance (pseudo R^2^ = 0%; RMSE = 27.6%), with high heterogeneity (*Ι* ^2^ = 93%; *Ι* ^2^_1_ = 89%; *Ι* ^2^_2_ = 3%). The presence of NDF concentration in models 2 and 3 (identical) improved model fit (RMSE = 25.4%; pseudo R^2^ = 21.9%), and slightly reduced total and between-trial heterogeneity (*Ι* ^2^ = 90%; *Ι* ^2^_1_ = 86%; *Ι* ^2^_2_ = 4%) compared to model 1. This model contained NDF concentration (*P* = 0.013), 3-NOP dose (*P* < 0.001) and predicted 35.0% reduction (intercept; *P* < 0.001) in RMD in CH_4_ yield when predictors were at their means. Similar to previous models, concentration of NDF was positively associated with CH_4_ output, predicting an 0.99 percentage point increase in RMD in CH_4_ yield for every 1 percentage point increase in NDF concentration. Model 4 identified OM concentration as a significant predictor (*P* = 0.020), in addition to 3-NOP dose (*P* < 0.001), which was negatively associated with CH_4_ yield. The model intercept estimated a 37.4% reduction (intercept; *P* < 0.001) when all moderators were at their mean. This model achieved the best overall fit (RMSE = 24.0%; pseudo R^2^ = 47.4%), with lower heterogeneity (*Ι* ^2^ = 84%; *Ι* ^2^_1_ = 80%; *Ι* ^2^_2_ = 4%) compared to models 1-3. Publication bias was evaluated by visual inspection of funnel plots (Supplementary Material, Figure S2) and by Egger’s regression tests. The standardized residuals for all CH_4_ outcomes showed no meaningful asymmetry, and Egger’s tests indicated no evidence of publication bias (*P* ≥ 0.05) across all models.

**Table 6.**
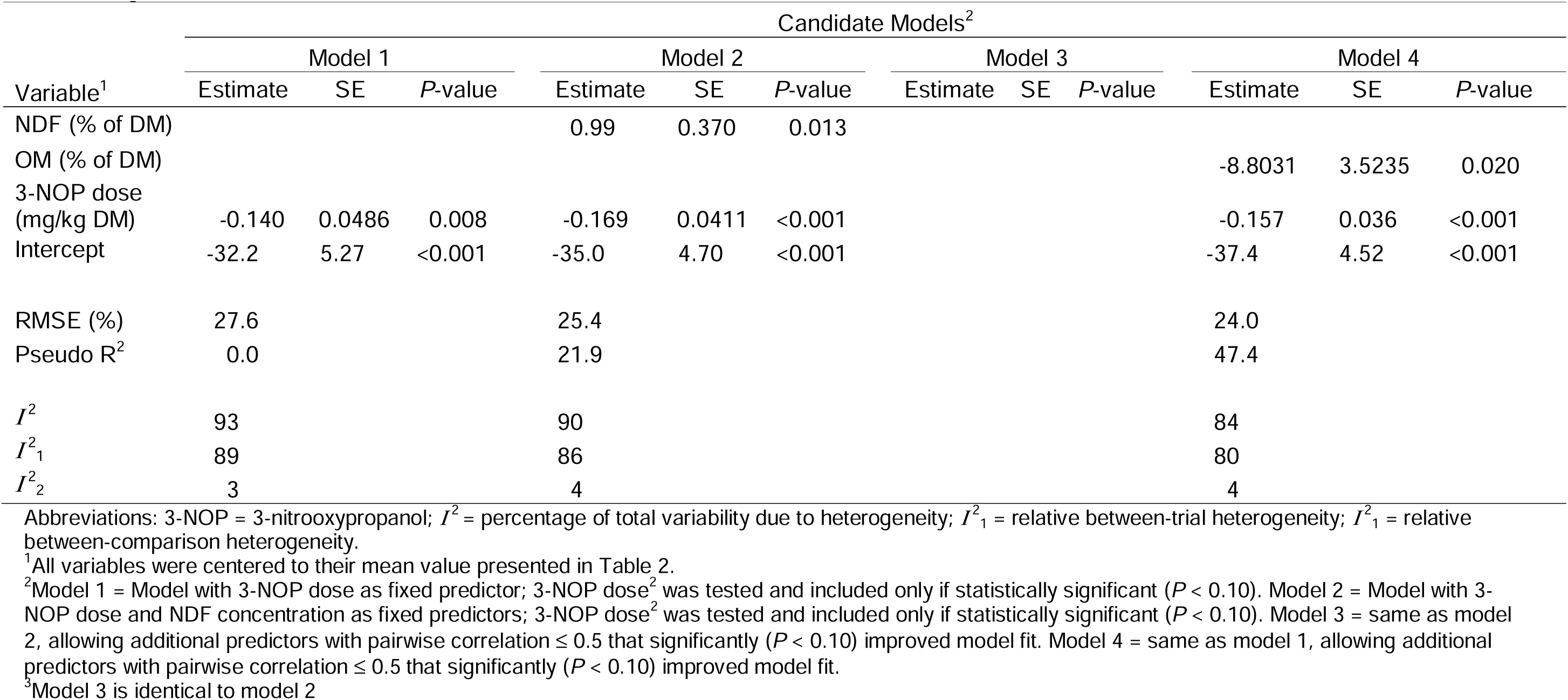
Estimates of overall 3-nitrooxypropanol (3-NOP) effect size and of explanatory variables from models for relative mean difference in methane yield (%)

## Discussion

The current meta-analysis represents the most comprehensive quantitative synthesis to date of 3-NOP effects in beef cattle, incorporating data from 17 studies across diverse production systems. Studies spanned from backgrounding to finishing production phases, and with 3-NOP dose ranging from 32 to 338 mg/kg DM. Despite variability in individual study responses, the results demonstrate consistent anti-methanogenic effect of 3-NOP in beef cattle production systems (Figure 2).

To characterize and predict this response, meta-regression models were developed for absolute CH_4_ production (g/d) and yield (g/kg of DMI), and for the relative change (%) in both metrics. Model 3 pre-included NDF concentration next to the intercept and 3-NOP dose, which was a key dietary factor influencing 3-NOP efficacy in other meta-analyses (Dijkstra et al., 2018; Kebreab et al., 2023). Additional uncorrelated variables that significantly improved model fit could also be included in model 3. This model, although occasionally yielding identical estimates to Models 2 or 4, offered the best balance of biological interpretability, statistical significance, predictive accuracy (RMSE), and was therefore selected as reference model. Hence, the subsequent discussion will be limited to model 3 for all CH_4_ metrics.

The antimethanogenic effects of 3-NOP supplementation on CH_4_ emissions in beef cattle, and the modifying roles of dietary NDF concentration (all CH_4_ expressions) BW, and DMI (the latter two only for CH_4_ production), are quantified through the predictive equations for each response variable (Model 3; Tables 3 to 6), as follows:

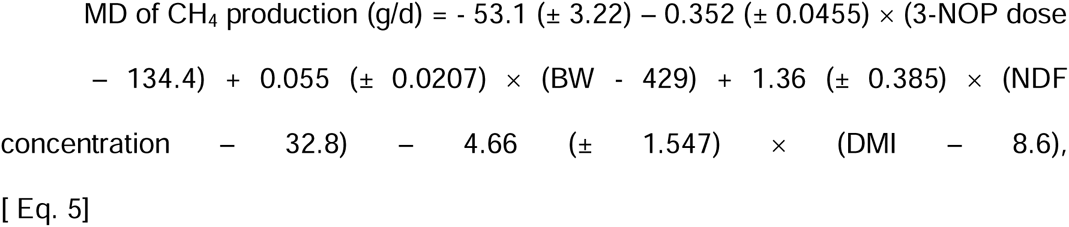

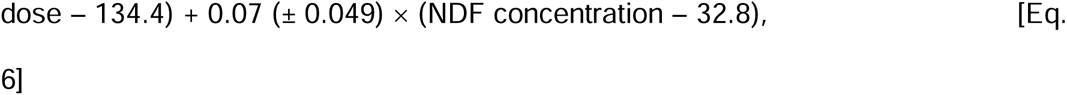

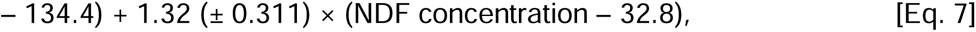

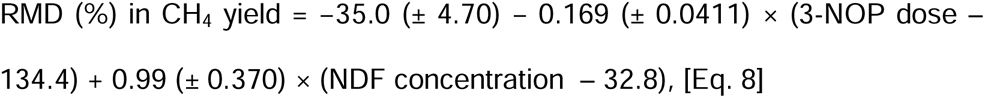

where, for all models, 3-NOP dose in mg/kg of DM, NDF concentration in % of DM, BW in kg, and DMI in kg/d.

At average predictor values, the MD equations predict absolute reductions of 53.1 g/d and 5.88 g/kg of DMI, whereas the RMD equations predict reductions of 37.6% in CH_4_ production and 35.0% in CH_4_ yield. These reductions are substantially greater than previously reported for beef cattle by Dijkstra et al. (2018), likely reflecting the greater number and diversity of studies included in the present meta-analysis (45 treatment means vs. 18 in earlier work), which allowed for a more robust and representative assessment of 3-NOP efficacy. In addition, average 3-NOP dose of the studies included in the Dijkstra et al. (2018) analyses was lower (i.e., 123 mg/kg DM). At this average dose, our model results in an average reduction of 35.2 and 33.1% in CH_4_ production and yield, respectively, which is substantially greater than 22.2 and 17.1% reported for beef cattle by Dijkstra et al. (2018). Compared to dairy systems, the observed mitigation efficacy exceeds the 32.7 and 30.9% reductions in CH_4_ production and yield, respectively, reported by Kebreab et al. (2023). This is most likely due to the lower average 3-NOP dose in dairy cattle (70.5 mg/kg DM) compared to beef cattle (134.4 mg/kg DM). At a dose of 70.5 mg/kg DMI the predicted relative reductions in CH_4_ production and yield in the current analysis are 24.2% for both units, being lower than the values reported for dairy cattle by Kebreab et al. (2023). This supports the conclusion of Dijkstra et al. (2018) that, at similar 3-NOP doses and NDF concentrations, the CH_4_ mitigating effect of 3-NOP is less pronounced in beef cattle than in dairy cattle.

Applying these MD models for the specific production phases in beef cattle, absolute reductions of 47.0 g/d and 6.3 g/kg of DMI for backgrounding (3-NOP dose = 163.4 mg/kg of DM; NDF concentration = 41.2% of DM, and DMI = 7.6 kg/d; Supplementary Material, Table S1), and 59.8 g/d and 5.4 g/kg of DMI for finishing cattle (3-NOP dose = 104.2 mg/kg of DM; NDF concentration = 24.0% of DM, and DMI = 9.7 kg/d; Supplementary Material, Table S1) are observed. For the RMD models, CH_4_ production reduced by 32.6% and 42.9%, and CH_4_ yield by 31.6% and 38.6%, in backgrounding and finishing cattle, respectively. Thus, although finishing cattle receive lower doses of 3-NOP, their lower NDF concentration and greater DMI contributes to greater absolute as well as relative reductions in CH_4_ production and yield, compared to backgrounding. Further combinations of different doses and NDF concentration along with the expected change in CH_4_ emissions are presented in the Supplementary Material, Table S2 and S3.

Dietary composition appears an essential modifier of 3-NOP efficacy, with dietary NDF concentration showing a particularly strong relationship with a reduction of 3-NOP efficacy. This finding aligns with mechanistic studies demonstrating that high-fiber diets promote hydrogen production through enhanced fibrolytic activity (van Lingen et al., 2021), potentially overriding 3-NOP’s capacity to redirect reducing equivalents, and leading to relatively less 3-NOP efficacy. This effect on CH_4_ production and yield (1.32 and 0.99 percentage point increase per 1 percentage point change in NDF concentration) appears more pronounced in beef cattle than the 0.63 percentage points impairment of 3-NOP efficacy per 1 percentage point change of NDF concentration established previously for dairy cattle (average NDF concentration 32.9% of DM; Kebreab et al., 2023). Our findings are in support of the hypothesis postulated in the study with beef cattle by Vyas et al. (2018) that structural carbohydrates may physically protect methanogens or alter rumen retention dynamics in ways that reduce methanogen exposure to 3-NOP.

Dry matter intake was identified as a significant predictor in models of MD CH_4_ production, with higher DMI associated with greater mitigation efficacy. It reflects the increased 3-NOP intake at higher DMI levels, where more additive would be ingested on a daily basis (mg/d). Greater DMI has been previously observed as an enhancer of 3-NOP efficacy when considered alone for models of RMD in CH_4_ intensity, but not CH_4_ production and yield, in dairy cattle (Kebreab et al., 2023). This effect may be particularly pertinent for beef cattle since they require larger 3-NOP doses compared to dairy cows to achieve similar mitigating effects (Dijkstra et al., 2018). Additionally, achieving effective doses also depends on the temporal pattern of intake and the physical form of delivery. For instance, confined beef cattle typically consume about 80% of the feed offered (as fed) within 12 h (Lee et al., 2015a; Lee et al., 2015b). Romero-Perez et al. (2015) observed a 59% CH_4_ reduction when 280 mg/kg DM of 3-NOP was mixed into a TMR, compared with a 37% reduction when the additive was top-dressed at 245 mg/kg DM in similar backgrounding diets, highlighting the influence of delivery methods in mitigation outcomes. Therefore, strategies to improve the consistency of 3-NOP exposure, such as more frequent feeding or slow-release formulations, may enhance mitigation potential in beef production systems.

Although the quadratic 3-NOP term was not identified in our selected models, it was retained in model 1 for MD in CH_4_ production. Its presence indicates that the effect of increasing dose on CH_4_ reduction appears saturable in the dose-response curve. This effect was not observed for RMD models, as observed in earlier meta-analyses (e.g., Kebreab et al., 2023). Our results reveal that while increasing the 3-NOP dose initially drives substantial reductions in CH_4_ production, diminishing returns emerge at 3-NOP levels exceeding ∼200 mg/kg of DM (Figure 3). This nonlinear pattern likely reflects multiple interacting constraints within the rumen environment. Mechanistically, it may arise from enzyme saturation and an adaptive microbial response. Duin et al. (2016) demonstrated that 3-NOP potently inactivates MCR at low micromolar concentrations. However, they also showed that MCR inhibition is reversible in vivo, due to the ability of methanogens to regenerate the active Ni(I) state of the enzyme via ATP- and hydrogen-dependent repair mechanisms. Thus, at high 3-NOP concentrations, the inhibition of methanogenesis may decline as methanogens either become saturated with the inhibitor or partially recover by increasing their enzymatic activity. The current findings have practical implications. From a feed management perspective, increasing 3-NOP doses beyond some 200 mg/kg of DM may yield marginal gains in CH_4_ abatement but with disproportionate increases in cost, especially considering the expense of precision delivery systems in feedlot settings.

Monensin was not retained as a predictor in any of the final selected models. Among the best candidate models ranked by RMSE across all four response variables, monensin appeared as a predictor in one subset of model 4 for RMD in CH_4_ production (*P* = 0.058; RMSE = 21.0%; Supplementary Material, Figure S3 and Table S4), indicating a trend of 13.1 percentage points lower reduction by 3-NOP when monensin is present in the diet. However, its inclusion failed to improve model accuracy and was therefore excluded from the final selected models. Herein, monensin was present in 8 of the 17 studies in the dataset, one of which specifically compared the individual and combined effects of 3-NOP and monensin in both backgrounding and finishing diet (Vyas et al., 2018). A significant interaction between 3-NOP and monensin on CH_4_ production was observed in beef cattle fed high forage diets. In qualitative agreement with the RMD model results (Table S4), the decrease in CH_4_ production due to 3-NOP was markedly smaller in the presence than in the absence of monensin. However, this differential effect did not occur when CH_4_ was expressed per unit of DMI. These results appear to reflect a numerical decrease in DMI with 3-NOP in the absence of monensin, and an unexpected numerical increase in DMI with 3-NOP in the presence of monensin. No significant interactions between 3-NOP and monensin were observed for CH_4_ production or yield in beef cattle fed high grain diets (Vyas et al., 2018). The absence of interaction for CH_4_ yield in both high forage and high grain diets may be attributed to the distinct mechanisms of action of these two feed additives. As previously described, 3-NOP directly inhibits the MCR enzyme, whereas monensin acts by inhibiting gram-positive bacteria and indirectly reduces hydrogen availability for methanogenesis (Russell and Houlihan, 2003). Thus, although monensin is commonly used in commercial feedlots diets to primarily improve animal performance and feed efficiency (Ribeiro et al., 2025), further studies directly including monensin as a treatment factor are necessary to disentangle the effect of diet composition and monensin on the efficacy of 3-NOP.

In the present analysis, RMD models showed larger RMSE than those reported for dairy cattle (Kebreab et al., 2023), indicating greater variability in relative reductions across beef studies. For instance, the RMD for CH_4_ production with dose and NDF as predictors (model 2 or 3; Table 5) had an RMSE of 23.3%, compared with the 7.34% in the equivalent dairy model (Kebreab et al., 2023). Similarly, for CH_4_ yield, RMSE is 25.4% (models 2 or 3; Table 6) versus 7.76% for dairy (Kebreab et al., 2023). This discrepancy arises partly from the wide range of baseline CH_4_ emissions in beef control groups (52 to 276 g/d), which amplifies variability when expressed as relative reductions.

Although this meta-analysis presents the most robust assessment to date on the effects of 3-NOP in beef cattle, this study has several limitations that should be considered when interpreting the results. One such limitation is the variation of 3-NOP supplementation duration among the included studies, which ranged from 3 to 34 weeks. As a result, the estimated effects may not fully reflect potential rumen adaptation on the longer term and for this purpose be biased towards observations in short-term studies. Long-term studies with beef cows would be valuable to assess whether shifts in rumen microbiome could reduce 3-NOP’s sustained effectiveness (Yu et al., 2021; Orzuna-Orzuna et al., 2024). Second, predictive models were developed using data within a specific dose range, from 32 to 338 mg/kg of DM, dietary NDF concentrations from 14.8% to 66.1% of DM, DMI from 5.7 to 11.8 kg/d, and BW from 147 to 732 kg. Applying these equations outside of these ranges may compromise their reliability and is therefore not recommended. Furthermore, at the extremes of the parameter space, such as very low 3-NOP doses combined with very high NDF concentrations, our models may predict minimal or even numerically positive changes in CH_4_ emissions relative to control. This observation reflects data points included in this meta-analysis (e.g., Vyas et al., 2018), highlighting the diet-dependent efficacy of 3-NOP and the importance of considering these specific parameter combinations. Third, the efficacy estimates rely on the assumption of proper mixing of 3-NOP and consistent intake by the animals, which can be difficult to achieve, especially in pasture-based systems where delivery is less controlled. For instance, in the only study included that evaluated tropical forage diets (Martínez-Fernández et al., 2018), 3-NOP was mixed with molasses and hay and offered at staggered intervals post-feeding (0, 3, and 6 hours) to prolong its presence in the rumen. This study reported promising reductions in CH_4_ production (30%) and CH_4_ yield (38%) at a 3-NOP dose of 338 mg/kg of DM. These values are lower than those predicted by our equations for production (36.4%), but greater for yield (36.4%) at the same NDF concentration of 66.1% of DM. Nonetheless, additional research is required to explore 3-NOP effects in tropical forage diets with different 3-NOP doses and delivery strategies. Fourth and last, the efficacy estimates in the present analysis are based largely on the assumption that the targeted or formulated 3-NOP dose reflects the actual dose consumed. However, deviations between targeted and realized doses can markedly affect CH_4_ mitigation outcomes due to the strong dose-dependent response observed in all derived equations. Studies that do not quantify actual 3-NOP intake may over- or underestimate its effectiveness, limiting comparability across studies and potentially informing mitigation policies inaccurately. For instance, van Gastelen et al. (2022) reported analyzed 3-NOP concentrations that were 7% lower and 9% higher, respectively, than targeted in grass silage- and corn silage-based diets fed to dairy cattle, which may explain the lower CH_4_ mitigation observed with grass silage in their experiment. Despite its importance, most studies do not report analyzed dietary 3-NOP levels in the diet, but report inclusion levels achieved based on amounts of 3-NOP mixed into the diet, perhaps also due to the uncertainty with collecting representative samples of mixed diets. As a result, our meta-analysis relied primarily on targeted doses, potentially introducing variability or bias. This misalignment may complicate the practical interpretation and application of the derived equations, especially in regulatory or farm-level settings. It is therefore recommended that future studies analyze and report the actual 3-NOP concentrations in feed after mixing and prior to feeding, and use analyzed rather than formulated dose when evaluating CH_4_ mitigation.

## Conclusions

The present meta-analysis confirms that 3-NOP is a highly effective feed additive for mitigating enteric CH_4_ emissions in beef cattle. On average, 3-NOP supplementation reduced CH_4_ production by 36.2 ± 24.43% and CH_4_ yield by 33.2 ± 25.54%. The developed models predicted reductions of 37.6% in CH_4_ production and 35.0% in CH_4_ yield when all predictors included in the model are at their mean value. Mitigation efficacy declined with increasing NDF concentration. These results support the effectiveness of 3-NOP in mitigating enteric CH_4_ emission in beef cattle and provide quantitative models to be used in assessment tools and GHG inventory methodology.

## Supplementary material

## Ethics approval

Not applicable.

## Data and model availability statement

The data/models that support the study findings are available upon request.

## Declaration of generative AI and AI-assisted technologies in the writing process

The authors did not use any artificial intelligence assisted technologies in the writing process.

## Acknowledgements

None.

