## Supplemental Tables and Figures for "A meta-analysis of 3-nitrooxypropanol effects on methane production and yield in beef cattle"

*animal* Journal

**Supplementary Material**

**Model description**

The mixed-effect meta-regression model was given by

$$y_{ij}=\beta_{0}+{\boldsymbol{x}_{ij}}^{T}\boldsymbol{\beta}+t_{i}+u_{j\left( i \right)}+e_{j\left( i \right)}$$

where $y_{ij}$ is the response mean difference ($\text{MD)}$ or relative mean difference $(\text{RMD}$) of the $j$^th^ treatment group comparison in the $i$^th^ publication, $\beta_{0}$ is the global intercept, ${\boldsymbol{x}_{ij}}^{T}$ is the row-vector of predictors corresponding to the $j$^th^ treatment group comparison in the $i$^th^ publication, $\boldsymbol{\beta}$ is the vector of fixed effects associated with the predictors, $t_{i}$ is the random effect of the $i$th publication, $u_{j\left( i \right)}$ is the random effect of the $j$th treatment group comparison nested in the $i$th publication and $e_{j\left( i \right)}$ is the sampling error with known variance calculated based on the standard error reported by the publication.

Supplementary Table 1: Descriptive statistics of feed intake, dietary characteristics, and methane emission by production phase^1^

|  | Backgrounding | | | | |  | Finishing | | | | |
| --- | --- | --- | --- | --- | --- | --- | --- | --- | --- | --- | --- |
| Variables | Mean | Median | SD | Minimum | Maximum |  | Mean | Median | SD | Minimum | Maximum |
| DMI (kg/d) | 7.6 | 7.0 | 1.68 | 5.7 | 11.7 |  | 9.7 | 10.1 | 1.41 | 6.9 | 11.8 |
| CP (% of DM) | 13.5 | 13.7 | 1.78 | 9.9 | 17.7 |  | 14.0 | 14.3 | 0.95 | 12.2 | 15.1 |
| NDF (% of DM) | 41.2 | 41.6 | 7.17 | 29.0 | 66.1 |  | 24.0 | 27.0 | 5.12 | 14.8 | 29.0 |
| Fat (% of DM) | 3.4 | 3.0 | 1.15 | 2.8 | 8.3 |  | 4.3 | 4.1 | 1.34 | 2.4 | 6.7 |
| Starch (% of DM) | 24.1 | 24.0 | 7.03 | 2.2 | 32.1 |  | 46.6 | 47.0 | 6.69 | 34.6 | 56.2 |
| OM (% of DM) | 92.6 | 92.4 | 0.68 | 91.0 | 93.6 |  | 95.1 | 95.4 | 1.26 | 92.3 | 96.6 |
| Roughage proportion (% of diet DM) | 70.1 | 65.0 | 12.19 | 50.0 | 100.0 |  | 9.3 | 8.0 | 6.38 | 3.1 | 27.6 |
| BW (kg) | 432.3 | 376.0 | 146.96 | 147.0 | 732.0 |  | 424.9 | 391.0 | 93.61 | 356.0 | 697.0 |
| Monensin dose (mg/kg DM) | 0.0011 | 0 | 0.00138 | 0.0 | 0.0033 |  | 0.0022 | 0.0026 | 0.00106 | 0 | 0.0033 |
| 3-NOP dose (mg/kg DM) | 163.4 | 150.0 | 73.11 | 39.6 | 337.8 |  | 104.2 | 100.0 | 43.4 | 32.0 | 200.0 |
| CH_4_ production (g/d) | 182.6 | 176.3 | 38.71 | 138.8 | 275.7 |  | 123.5 | 124.5 | 42.27 | 51.8 | 204.5 |
| MD CH_4_ production (g/d) | -45.8 | -45.0 | 28.17 | -102.9 | 10.2 |  | -54.2 | -55.8 | 29.07 | -97.8 | 3.6 |
| Relative MD CH_4_ production (% of control) | -25.0 | -28.0 | 15.39 | -59.2 | 7.1 |  | -48.0 | -47.5 | 26.80 | -90.6 | 2.9 |
| CH_4_ yield (g/kg DMI) | 24.5 | 23.8 | 2.14 | 19.7 | 28.6 |  | 12.8 | 13.9 | 4.30 | 5.1 | 19.1 |
| MD CH_4_ yield (g/kg DMI) | -5.8 | -5.4 | 3.81 | -14.0 | 1.8 |  | -4.8 | -4.7 | 3.41 | -13.0 | 3.2 |
| Relative MD CH_4_ yield (% of control) | -23.6 | -21.6 | 15.20 | -59.1 | 7.6 |  | -43.3 | -42.2 | 30.27 | -88.8 | 22.4 |

Abbreviations: DMI = DM intake; OM = Organic matter; 3-NOP = 3-nitrooxypropanol; CH_4_ = Methane

^1^Summaries for all variables reflect the full dataset (including control and 3-NOP treatments), except for CH_4_ production and CH_4_ yield, which are presented as control group means only. Mean difference (MD) values relate to 3-nitrooxypropanol (3-NOP) treatment mean compared with the control treatment mean.

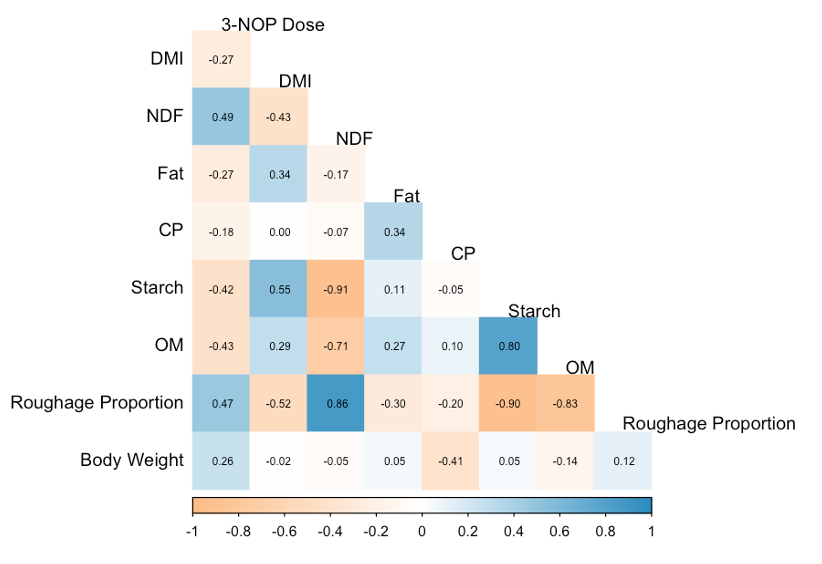

Supplementary Figure 1. Correlations of the explanatory variables considered in the meta-regression. Abbreviations: DMI = DM intake; 3-NOP = 3-nitrooxypropanol; OM = Organic matter.

**Supplementary Table 2**

Model-based expected relative mean difference (MD) in CH_4_ production (g/d) and yield (g/kg DMI) for indicative values of NDF and doses of 3-nitrooxypropanol (3-NOP) for backgrounding cattle.

| Backgrounding | | | |
| --- | --- | --- | --- |
| NDF (% DM) | Dose (mg/kg DM) | Change in CH_4_ production (%) | Change in CH_4_ yield (%) |
| 30 | 40 | -21.5 | -21.8 |
| 30 | 100 | -34.1 | -32.0 |
| 30 | 150 | -44.6 | -40.4 |
| 30 | 200 | -55.1 | -48.9 |
| 30 | 250 | -65.6 | -57.3 |
| 30 | 338 | -84.1 | -72.2 |
| 40 | 40 | -8.3 | -11.9 |
| 40 | 100 | -20.9 | -22.1 |
| 40 | 150 | -31.4 | -30.5 |
| 40 | 200 | -41.9 | -39.0 |
| 40 | 250 | -52.4 | -47.4 |
| 40 | 338 | -70.9 | -62.3 |
| 50 | 40 | 4.9 | -2.0 |
| 50 | 100 | -7.7 | -12.2 |
| 50 | 150 | -18.2 | -20.6 |
| 50 | 200 | -28.7 | -29.1 |
| 50 | 250 | -39.2 | -37.5 |
| 50 | 338 | -57.7 | -52.4 |
| 60 | 40 | 18.1 | 7.9 |
| 60 | 100 | 5.5 | -2.3 |
| 60 | 150 | -5.0 | -10.7 |
| 60 | 200 | -15.5 | -19.2 |
| 60 | 250 | -26.0 | -27.6 |
| 60 | 338 | -44.5 | -42.5 |
| 65 | 40 | 24.7 | 12.8 |
| 65 | 100 | 12.1 | 2.7 |
| 65 | 150 | 1.6 | -5.8 |
| 65 | 200 | -8.9 | -14.2 |
| 65 | 250 | -19.4 | -22.7 |
| 65 | 338 | -37.9 | -37.5 |

Abbreviations: DMI = DM intake; CH_4_ = Methane.

**Supplementary Table 3**

Model-based expected relative mean difference (MD) in CH_4_ production (g/d) and yield (g/kg DMI) for indicative values of NDF and doses of 3-nitrooxypropanol (3-NOP) for finishing cattle.

| Finishing | | | |
| --- | --- | --- | --- |
| NDF (% DM) | Dose (mg/kg DM) | Change in CH4 production (%) | Change in CH4 yield (%) |
| 15 | 32 | -39.6 | -35.3 |
| 15 | 100 | -53.9 | -46.8 |
| 15 | 125 | -59.1 | -51.0 |
| 15 | 150 | -64.4 | -55.3 |
| 15 | 175 | -69.6 | -59.5 |
| 15 | 200 | -74.9 | -63.7 |
| 20 | 32 | -33.0 | -30.4 |
| 20 | 100 | -47.3 | -41.9 |
| 20 | 125 | -52.5 | -46.1 |
| 20 | 150 | -57.8 | -50.3 |
| 20 | 175 | -63.0 | -54.5 |
| 20 | 200 | -68.3 | -58.8 |
| 25 | 32 | -26.4 | -25.4 |
| 25 | 100 | -40.7 | -36.9 |
| 25 | 125 | -45.9 | -41.1 |
| 25 | 150 | -51.2 | -45.4 |
| 25 | 175 | -56.4 | -49.6 |
| 25 | 200 | -61.7 | -53.8 |
| 30 | 32 | -19.8 | -20.5 |
| 30 | 100 | -34.1 | -32.0 |
| 30 | 125 | -39.3 | -36.2 |
| 30 | 150 | -44.6 | -40.4 |
| 30 | 175 | -49.8 | -44.6 |
| 30 | 200 | -55.1 | -48.9 |

Abbreviations: DMI = DM intake; CH_4_ = Methane.

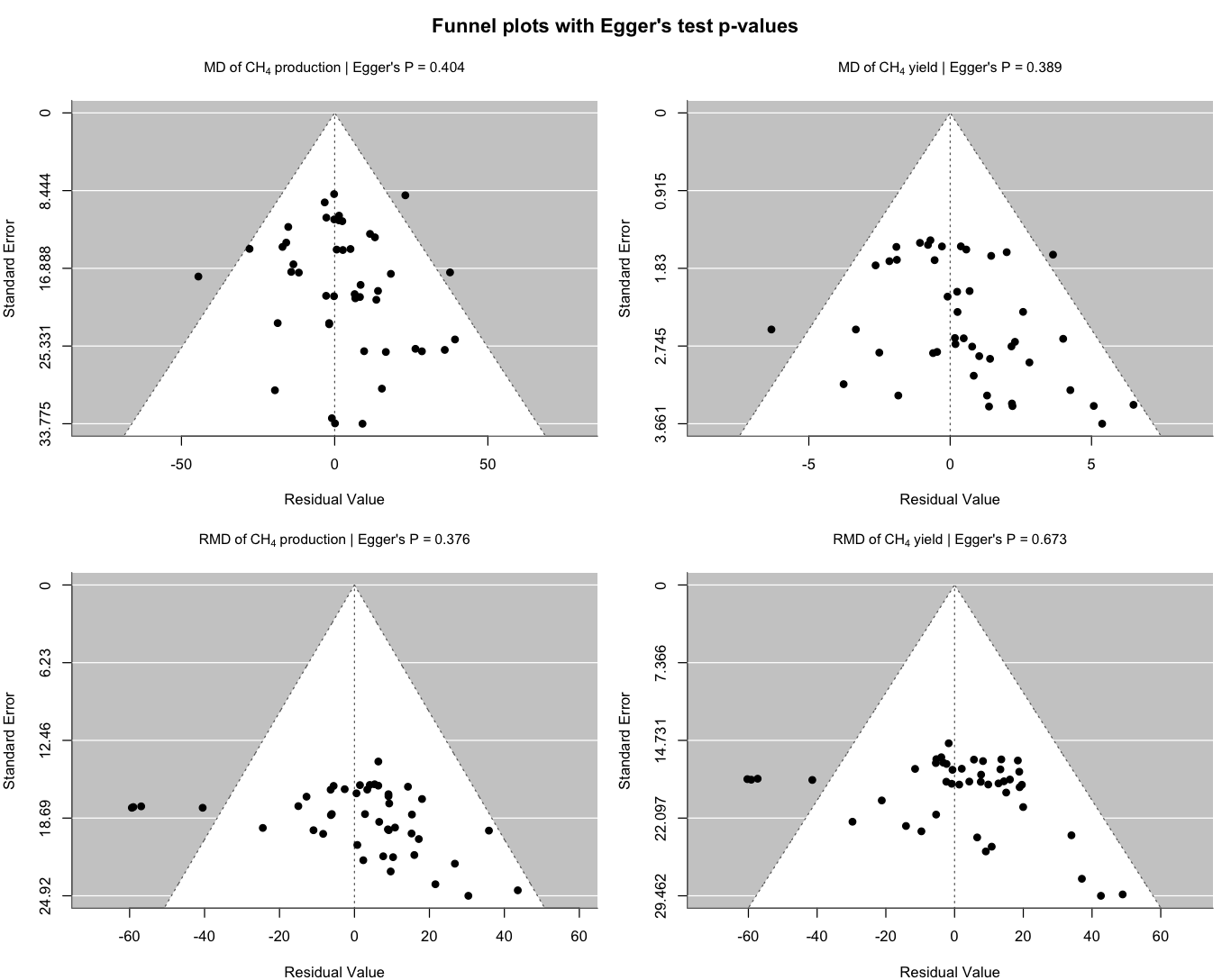

**Supplementary Figure 2.** Funnel plots of standardized residuals for mean difference (top row) and relative mean difference (bottom row) of CH_4_ production (g/d) yield (g/kg of DMI). Egger’s test *P*-values indicate no significant asymmetry (*P* ≥ 0.05).

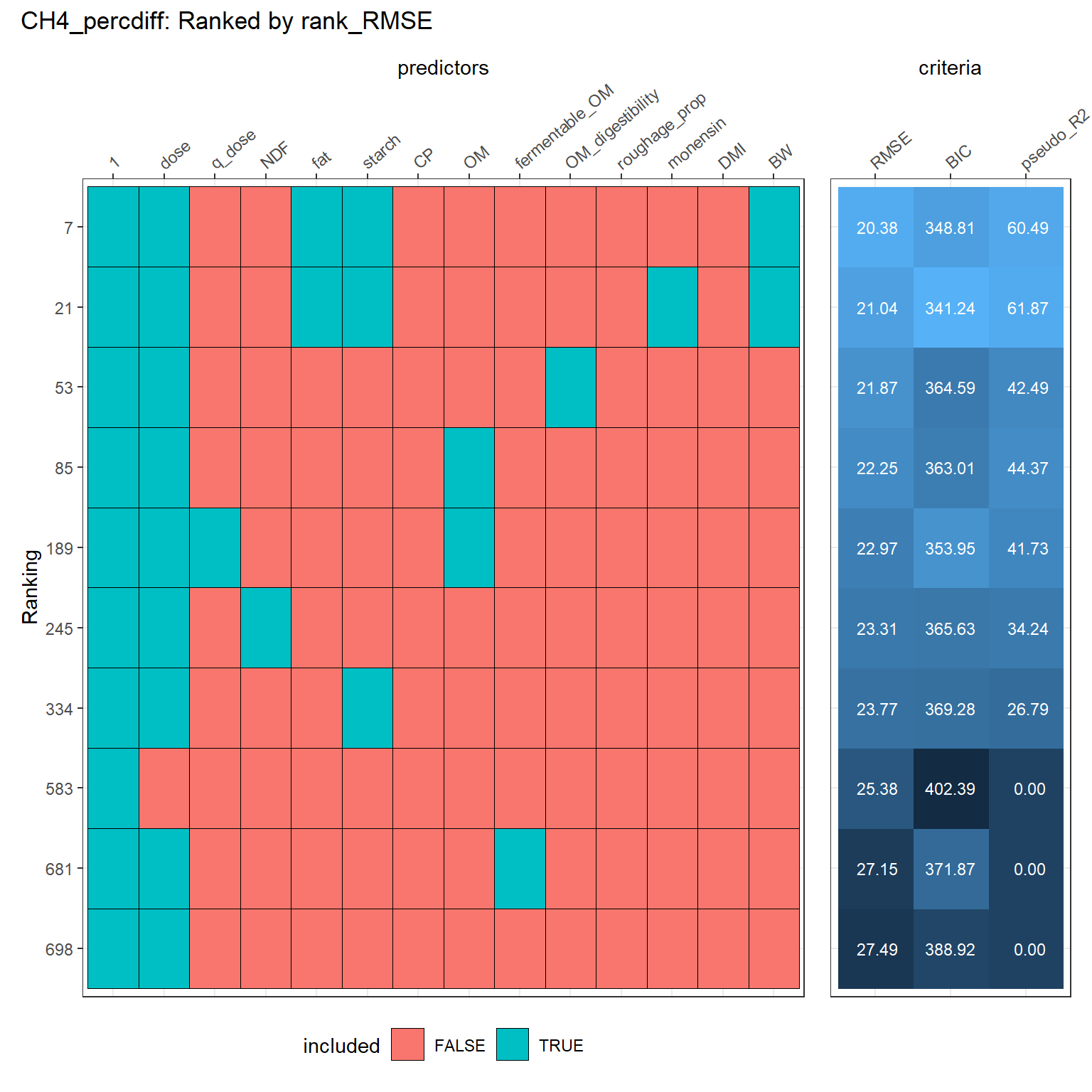

**Supplementary Figure 3.** Subset of models (Model 4) for relative mean difference in CH_4_ production (g/d). Each row represents a different model ranked by performance (RMSE), and each column shows whether a given predictor was included (blue) or excluded (red) in that model. Abbreviations: DMI = DM intake; OM = Organic matter; BIC = Bayesian information criterion.

**Supplementary Table 4**

Estimates of overall 3-nitrooxypropanol (3-NOP) effect size and of explanatory variables of relative mean difference in methane production (%), selected from candidate models ranked by RMSE (Ranking 21; Supplementary Figure 3).

|  | Model 598 | | |
| --- | --- | --- | --- |
| Variable | Estimate^1^ | SE | *P*-value |
| Fat (% of DM) | -7.77 | 1.980 | <0.001 |
| Starch (% of DM) | -1.19 | 0.197 | <0.001 |
| BW (kg) | 0.06 | 0.023 | 0.101 |
| 3-NOP dose | -0.21 | 0.032 | <0.001 |
| Monensin inclusion | 13.1 | 6.70 | 0.058 |
| Intercept | 29.9 | 12.66 | 0.057 |
| RMSE (%) | 21.0 |  |  |
| Pseudo R^2^ | 61.9 |  |  |

Abbreviations: 3-NOP = 3-nitrooxypropanol.

^1^Estimates are not centered at the mean values.
